# Lethal epistasis maintains strong linkage disequilibrium between unlinked supergenes

**DOI:** 10.64898/2026.01.08.698410

**Authors:** Giulia Scarparo, Azariah Lopez, Elijah Muro, Alan Brelsford, Jessica Purcell

## Abstract

Linkage disequilibrium (LD) between adaptive gene combinations is typically maintained through physical linkage and suppressed recombination. Although epistatic interactions can maintain LD between unlinked loci, this mechanism is rarely documented in nature. Supergenes, regions of suppressed recombination containing tightly linked loci, typically exemplify the first mechanism, in which genomic rearrangements lock together coadapted alleles. Here, we demonstrate a rare case of epistasis-driven LD between two supergenes on different chromosomes in the European ant *Formica cinerea*: one controlling colony queen number (chromosome 3) and another determining sexual body size (chromosome 9). We show that these supergenes assort independently according to Mendelian expectations during meiosis, yet exhibit high LD between the multi-queen haplotype P_2_ and the miniaturizing haplotype 9r, with small queens and males occurring exclusively in multi-queen colonies. Mismatched genotype combinations (P_2_ without 9r and vice versa) are severely underrepresented among all adult castes, with mismatched males experiencing complete mortality. We show that this pattern cannot be explained by meiotic drive or maternal-effect killing, assortative mating, or extrinsic selection on adults, indicating strong postzygotic epistatic selection maintaining the observed LD. The fitness costs of these epistatic interactions are substantial but critically depend on mating combinations: heterozygous small queens achieve 1.5-2.5 times higher fitness (based on offspring genotype viability) when mated to small males (P_2_-9r) compared to large males. Our findings provide empirical evidence for epistatic interactions between unlinked supergenes maintaining LD in the absence of physical linkage.

## Introduction

Understanding the link between genotype, phenotype, and fitness is a key step toward unveiling the genetic mechanisms that drive evolutionary adaptation within species. In 1930, Fisher conceptualized adaptation as populations evolving toward a single fitness optimum through phenotypic changes caused by mutations, assuming a relatively smooth fitness landscape where mutations have predictable, consistent effects on fitness (Fisher 1930; reviewed by Orr 2005). In this regard, a mutation could be classified as beneficial or deleterious based on its intrinsic properties. Around the same time, Wright suggested a more complex view of the adaptive process, emphasizing the role of epistasis in shaping fitness (Wright 1931; reviewed by Bank 2022). Epistasis, defined as interactions between genes, creates rugged fitness landscapes with multiple adaptive peaks that correspond to different advantageous genetic combinations within a population.

In scenarios involving epistasis, the effect of a mutation on fitness is not intrinsic but depends on the genetic background in which it occurs. For example, an identical mutation can cause degenerative symptoms in one person but not in another (healthy carrier), depending on the genetic context (reviewed by Kammenga 2017). Epistatic interactions that influence survival are expected to produce some degree of linkage disequilibrium (LD), since individuals with high-fitness combinations of alleles will be more common than those with other combinations. In extreme cases, epistatic interactions can be fatal; Dobzhansky (1946) called this “synthetic lethality.” In *Drosophila pseudoobscura*, Dobzhansky (1946) inferred that recombination between two wild-derived haplotypes on the same chromosome caused complete mortality in a homozygous state. When the alternative haplotypes did not recombine, neither imposed a fitness cost on the offspring. Dobzhansky (1946) stated that *de novo* mutations could not be ruled out as the cause of the lethality in this context. Observing the same synthetic lethality in multiple independent recombination events between the same pair of loci would resolve this uncertainty. Some attempts have been made to model the expected frequency of such synthetic lethals in natural populations (Phillips & Johnson 1998), but empirical data documenting them remain scarce.

When the fitness consequences of epistatic interactions are severe, selection is expected to favor recombination modifiers that further increase LD under some conditions (Charlesworth 1993; Feldman et al. 1997). Broadening our understanding of the conditions that result in fitness-based LD *versus* the appearance of recombination modifiers in empirical systems is a much-needed step in exploring the evolutionary consequences of epistasis.

In cases with fitness-based LD, selection against maladaptive gene combinations can maintain linkage disequilibrium between beneficial allelic combinations, even without physical linkage. This creates functional associations that persist despite ongoing recombination. Selection maintaining epistatic interactions between physically unlinked loci has been documented across diverse organisms throughout the tree of life, with varying evolutionary consequences. For example, in the protozoan parasite *Toxoplasma gondii*, epistatic interactions among seven loci influence virulence traits (Lehmann et al. 2004). In wheat, epistasis between loci on different chromosomes may reduce the effective population size of crosses carrying incompatible allelic combinations (Tessele et al. 2025). In stick insects, epistatic interactions shape color morphs that experience differential survival (Nosil et al. 2020). However, if deleterious combinations of alleles are lethal, this imposes substantial fitness costs through the loss of nonviable offspring.

This high cost can be mitigated through the evolution of recombination suppression between interacting loci, which can be achieved through genome structural rearrangements, including inversions (if the loci are already on the same chromosome), translocations, or chromosomal fusions (Charlesworth & Charlesworth 1973). Classical models of sex chromosome evolution exemplify this mechanism. Suppressed recombination is favored when a sex-determining gene is linked to sexually antagonistic alleles, variants that are advantageous in one sex but detrimental in the other (Charlesworth and Charlesworth 1978; Charlesworth et al. 2005). This suppression of recombination can continue to expand in a stepwise fashion. Several empirical cases demonstrate how regions of suppressed recombination in ancestral sex chromosomes expand through chromosomal fusions with autosomes, creating neo-sex chromosomes. Examples include threespine sticklebacks (Kitano et al. 2009), various bird species (Gan et al. 2019; Huang et al. 2022, Muirhead et al. 2025), and butterflies (Mongue et al. 2017), although many of these examples do not clearly document a role of fitness epistasis in the evolution of neo-sex chromosomes.

Supergenes, clusters of tightly linked functional genetic elements that control discrete, complex phenotypes, typically exemplify the first mechanism, where genomic rearrangements lock together coadapted combinations of loci as evidence of past epistatic events (Gutiérrez-Valencia et al. 2021). However, epistatic interactions can still be detected within supergenes, as demonstrated in the land snail *Cepaea nemoralis*, where, despite suppressed recombination, some individuals exhibit intermediate coloration and banding patterns that deviate from the expected discrete morph classes (Gonzalez et al. 2019). At a larger scale, entire inversions can interact epistatically with each other, creating complex multi-locus systems where different chromosomal rearrangements function as interacting units. This phenomenon is observed in *Drosophila pseudoobscura*, where epistasis acts between three non-overlapping inversions on the same chromosome that together form a sex-ratio supergene (Fuller et al. 2020).

Despite the growing body of literature describing new supergenes in a variety of organisms (reviewed by Purcell & Brelsford 2025, and Gutiérrez-Valencia et al. 2021), we still know little about the role of epistasis in generating, maintaining, or expanding LD between loci. One promising avenue is to leverage natural systems in which multiple supergenes co-occur within a single species, enabling direct study of inter-supergene epistatic interactions. The recent discovery of an independently evolved two-supergene system in the ant *Temnothorax rugatulus* (Darras et al. 2026), together with the presence of multiple supergenes in the Atlantic cod (Matschiner et al. 2022), suggests that complex genetic architectures involving multiple supergenes may be more widespread than previously appreciated. This raises the possibility that interactions among supergenes represent a recurring feature of genome evolution and highlights the study of inter-supergene interactions as an emerging and timely research direction. Here, we take advantage of a rare natural system where two supergenes located on different chromosomes and associated with complex phenotypic traits co-occur within an ant species (Scarparo et al. 2023), providing an opportunity to investigate their interactions.

In multiple ant genera (*Solenopsis* (Wang et al. 2013), *Formica* (Purcell et al. 2014), *Pogonomyrmex* (Errbii et al. 2024), *Cataglyphis* (Lajmi et al. 2026), *Leptothorax* (Braim 2015), *Myrmica* (Sigeman et al. 2025a), *Myrmecina* (Mona et al. 2025), and *Temnothorax* (Darras et al. 2026)), supergenes have convergently evolved to control colony social form and other fitness-related traits (Purcell and Brelsford 2025, De Gasperin et al. 2024; Lawson et al. 2012; Scarparo et al. 2026). In *Formica*, an ancient supergene, dating back approximately 30 million years, is widespread within the genus and spans much of chromosome 3 (Brelsford et al. 2020; Purcell et al. 2021). In several *Formica* species, this supergene has been consistently linked to colony queen number (Purcell et al. 2014; McGuire et al. 2022; Pierce et al. 2022; Purcell et al. 2025), although exceptions have been observed (Scarparo et al. 2024; Lagunas-Robles et al. 2025; Sigeman et al. 2025b).

Recently, Scarparo et al. (2023) described a novel supergene system in the European *Formica cinerea*. This species carries four supergene haplotypes (M_A_, M_D_, P_1_, and P_2_) on chromosome 3, as well as a second supergene on chromosome 9 with two alternative haplotypes (9a and 9r). Beyond influencing colony queen number, this complex system is also associated with colony split sex ratio, similar to the results reported in several North American *Formica* species (Lagunas-Robles et al. 2021; Lagunas-Robles et al. 2026), and with alate body size. In *F. cinerea*, Scarparo et al. (2023) identified three distinct queen morphs (Table 1): (1) monogyne macrogyne queens, carrying only M haplotypes on chromosome 3 and homozygous 9a9a on chromosome 9; (2) polygyne macrogyne queens, approximately 5% smaller than their monogyne counterparts, carrying at least one P_1_ haplotype (but not P_2_) on chromosome 3 and 9a9a on chromosome 9; and (3) polygyne microgyne queens, up to 20% smaller than macrogyne queens, carrying at least one P_2_ haplotype on chromosome 3 and at least one 9r on chromosome 9. Males, that are haploid, carrying the P_2_–9r combination are, on average, 9% smaller than males without 9r (Table 1). In this initial study, the P_2_ haplotype showed strong LD with the 9r haplotype (Scarparo et al. 2023). The male and queen morphs in *F. cinerea* have been also recently associated with distinct dispersal, mating patterns and colony foundation strategies (Scarparo et al. 2026).

**Table 1.** Overview of *F. cinerea* supergene system. The supergene on chromosome 3 is associated with monogyny (the presence within a colony of a single reproductive queen) and polygyny (the presence of two or more reproductive queens). The supergene on chromosome 9 is associated with sexual body size, where large queens and males are respectively called macrogynes and macraners, and small queens and males are called microgynes and micraners.

| Genotype chr3 | Genotype chr9 | Colony social form | Sexual body size | Notes |
| --- | --- | --- | --- | --- |
| M<sub>A</sub>M<sub>A</sub><br>M<sub>A</sub>M<sub>D</sub><br>M<sub>D</sub>M<sub>D</sub> | 9a9a | Monogyne<br>(=single queen colony) | Macrogynes<br>(=large queens)<br>and macraners<br>(=large males) |  |
| M<sub>A</sub>P<sub>1</sub><br>M<sub>D</sub>P<sub>1</sub><br>P<sub>1</sub>P<sub>1</sub> | 9a9a | Polygyne (=multi queen colony) | macrogynes and macraners | polygyne macrogynes are 5% smaller than monogyne macrogynes; polygyne macraners are 2.6% bigger than monogyne macraners |
| M<sub>A</sub>P<sub>2</sub><br>M<sub>D</sub>P<sub>2</sub><br>P<sub>1</sub>P<sub>2</sub><br>P<sub>2</sub>P<sub>2</sub> | 9a9r<br>9r9r | Polygyne | Microgynes (=small queens) and micraners (=small males) | microgynes are up to 20% smaller than macrogynes; micraners are ~9% smaller than macraners |

Scarparo et al. (2023) proposed that small queens are strongly disadvantaged in a monogyne background, where they would be the sole reproductive individual, and that strong selection, therefore, could have favored linkage between the polygyne-associated haplotype P_2_ and the miniaturizing haplotype 9r.

Here, we investigate the nature of the observed LD between the two *F. cinerea* supergenes and ask whether the fitness effects of one supergene depend on the genetic state of another. We considered two possible mechanisms that could explain strong LD between P_2_ and 9r: physical linkage (e.g., chromosomal fusion or reciprocal translocation) that prevents recombination, or epistatic selection against mismatched genotypes (i.e., individuals carrying P_2_ without 9r, or 9r without P_2_) that maintains LD despite recombination. The occasional occurrence of mismatched genotypes suggests that individuals with mismatches have reduced survival, but that the genotypic mismatch is not completely lethal. This opens the possibility that genotype combinations between chromosomes 3 and 9 generate a complex fitness landscape with differential survival of various genotype combinations. To disentangle these mechanisms, we use three complementary approaches. First, we assess whether P_2_ and 9r exhibit gene drive, as supergenes are known to harbor selfish genetic elements that can bias their own transmission at the expense of non-selfish haplotypes (Chapuisat & Purcell 2026). Such transmission distortion could skew Mendelian expectations and contribute to the low frequency of mismatched genotypes observed in adults. Second, we characterize the pattern of LD between the two supergenes by analyzing genotype frequencies in eggs to test for independent assortment, which would occur under epistatic selection but not under physical linkage. Finally, we analyze genotype frequencies across sexes and morphological castes. Specifically, we compare expected and observed haplotype combinations in adults (workers, gynes, and males) using data from Scarparo et al. 2023, together with newly collected mated queens and spermathecal contents from this study, to detect deviations from expected haplotype combination frequencies. We further compare frequencies between reproductive individuals before and after mating (unmated gynes vs. mated queens; unmated males vs. spermathecal contents) to rule out deviations arising from mating failure rather than viability selection.

## Materials and Methods

### Sample collection

*Formica cinerea* queens were collected in the Aosta Valley and Piedmont regions in Italy between late June and early July over multiple years from 2019 to 2022. Queens were either collected directly from colonies along with several workers (classified as mature queens) or captured after mating flights while clinging to vegetation or walking on the ground in search of suitable nest sites (classified as newly mated queens). After collection, each queen was kept individually in a glass tube. Newly mated queens were provided only water, while mature queens were given a sugar-water solution and accompanied by 3–4 workers from their original colony. Brood production was monitored every other day, and eggs or larvae were collected once at least 10 brood items were present (this was repeated if queens continued to lay eggs). Queens were kept alive for an average of two months before being preserved in 100% ethanol for genetic analysis. Colony fragments with gynes (unmated queens), males and workers were collected from the same locations, during the same years and seasons, as the queens. These samples were previously sequenced and discussed in Scarparo et al. (2023) to describe the supergene system in *F. cinerea*.

### PCR-RLFP assays

We developed five PCR–restriction fragment length polymorphism (PCR-RFLP) assays to distinguish the four supergene haplotypes on chromosome 3, and two additional assays to differentiate the two haplotypes on chromosome 9 in *F. cinerea*. Each assay used two pairs of primers (see Table S1, columns “External primer sequences” and “Internal primer sequences”) to perform nested PCR, which is particularly effective when working with low DNA quantities, such as those extracted from single eggs or spermathecae. Each primer set was paired with a specific restriction enzyme targeting a diagnostic single-nucleotide polymorphism (SNP), which allowed us to distinguish haplotypes based on the presence or absence of restriction cuts in a short amplified fragment. For details on the DNA extraction and genotyping protocols, primer sequences, and corresponding enzymes, see the supplementary files, Table S1 and Figure S1. The complete dataset of all genotyped individuals through PCR-RFLP is provided in Table S2.

### Caveat

This study investigates the relationship between the P_2_ haplotype on chromosome 3 and the 9r haplotype on chromosome 9. Because both haplotypes occur exclusively in polygyne colonies (Scarparo et al. 2023), and given the complexity of chromosome 3, which harbors four alternative haplotypes, we restricted all analyses to a polygyne background to avoid reducing statistical power. This also avoids confounding effects arising from differences between monogyne and polygyne systems. For transmission analyses (gene drive and independent assortment), we further restricted the dataset to queens carrying exclusively P haplotypes (P_1_P_2_), excluding queens carrying the monogyne-associated M haplotype (MP_2_ queens). The interaction between P_2_ and 9r can be fully assessed using P_1_P_2_ queens alone; including MP_2_ queens would introduce additional complexity due to the presence of M haplotypes without providing further insight into this specific relationship. Although analyses comparing P_2_ vs. M (as well as P_1_ vs. M) would be valuable for understanding the maintenance of genetic polymorphism in the population, these are beyond the scope of the present study. Preliminary results for MP_2_ queens are provided in the Supplementary Material (Figure S2). Importantly, this filtering was applied only to the transmitting individuals (queens), while offspring were retained regardless of carrying an M haplotype (e.g., MP_1_ and MP_2_), as these arise through paternal inheritance. The dataset containing all individuals effectively analyzed in this study is provided in Table S3.

### Meiotic Drive and independent assortment of Chr 3 and 9 in eggs

In a previous study, Scarparo et al. (2023) observed strong association between the P_2_ and 9r haplotypes. However, the nature of this association remains unclear. If it is due to physical linkage, maternal P_2_ and maternal 9r should be co-transmitted nearly 100% of the time in offspring of queens heterozygous for both supergene haplotypes, with only rare exceptions caused by recombination events. Alternatively, if the association reflects selection against “mismatched” combinations (epistasis) later in development, we would expect the four possible maternal haplotype combinations (P_2_–9r, P_2_–9a, P_1_–9r, P_1_–9a; where the queen is P_1_P_2_-9a9r) to occur at equal frequencies (25% each) under independent assortment. However, non-random associations between haplotypes could also arise from biased transmission during meiosis or early in development (e.g. Keais et al. 2025). In particular, if one or both haplotypes is a meiotic driver, this could generate deviations from Mendelian expectations and contribute to the observed association between P_2_ and 9r independently of physical linkage or epistasis.

We first tested whether P_2_ on chromosome 3, 9r on chromosome 9, or both deviate from Mendelian segregation by genotyping 104 eggs from 12 P_1_P_2_-queens and 164 eggs from 20 9a9r queens. We then examined the association between maternal haplotypes on chromosomes 3 and 9 by genotyping 79 eggs from 10 P_1_P_2_–9a9r queens. Since diploid eggs receive one chromosome from each parent, maternal haplotypes transmitted to eggs were inferred by subtracting the known paternal contribution (determined by genotyping sperm from the spermatheca) from each egg’s diploid genotype. For example, if an P_1_P_2_ queen mated with an P_1_ male and produced 50% P_1_P_1_ eggs and 50% P_1_P_2_ eggs, the maternal haplotype transmission would be 50% P_1_ and 50% P_2_.

### Maternal-effect killing

In later developmental stages (e.g. larvae), haplotype frequencies could be distorted by gene drive mechanisms other than meiotic drive, such as the maternal-effect killer described in *F. selysi* (Avril et al. 2020). While Avril et al. (2020) focused on interactions between M and P haplotypes in *F. selysi*, here we investigate potential skews in genotype frequencies between eggs and larvae with respect to the P_1_ versus P_2_ haplotypes on chromosome 3 and the 9a versus 9r haplotypes on chromosome 9.

We calculated genotype frequencies for chromosomes 3 and 9 in eggs and larvae whose mothers are P_1_P_2_ (n=8 for 49 eggs and n=6 for 48 larvae) or 9a9r (n=13 for 78 eggs and n=11 for 81 larvae) and mated with X-9a males (M_A_-9a or P_1_-9a). We analyzed whether the proportion of P_2_ heterozygous and 9a9r offspring differed between eggs and larvae. Moreover, we investigated whether transmission of the four possible maternal haplotype combinations (P_2_–9r, P_2_–9a, P_1_–9r,P_1_–9a) from double heterozygous P_1_P_2_-9a9r queens occurred at equal frequencies in eggs and larvae (at the expected rate of 25% for each haplotype combination), or whether some combinations were underrepresented in larvae compared to eggs, indicating early mortality.

### Epistasis in adults

To test for signatures of epistasis that might have led to LD between P_2_ and 9r, we re-analyzed the dataset from Scarparo et al. (2023). In that study, Scarparo and cohautors collected *F. cinerea* colony fragments without prior knowledge of their genotypes and genotyped up to five workers, eight gynes, and eight males per colony using RAD-seq. Gynes and males were collected before mating flights, often during the pupal stage, and genotyped after eclosion. Importantly, the genotypes of the reproductive queens were not directly known. For the present study, we subset the original dataset from Scarparo et al. (2023) to include only individuals from polygyne colonies, for which social structure was inferred through relatedness analyses of at least five diploid individuals. We further excluded individuals from four colonies (n=6) that exclusively carried M haplotypes, representing exceptions to the general association between MM genotypes and monogyne colonies, and between genotypes carrying at least one P haplotype and polygyne colonies. These exceptions have previously been suggested to reflect declining monogyne colonies that may be more prone to accepting additional queens (Scarparo et al. 2023). In three additional colonies, one caste carried the MM genotype while another caste carried genotypes with at least one P haplotype; these colonies were retained in the dataset. The final dataset comprised 261 workers, 81 gynes, 135 haploid males. We also included an additional 62 mature queens carrying at least one P haplotype (23 analysed in Scarparo et al. 2023 and 39 collected in 2022 and reported here for the first time), as well as 40 of their spermathecae (Table S4).

We compared observed and expected genotype combinations separately for workers, gynes, mated queens, spermathecae, and males to assess whether certain combinations occurred more or less frequently than expected. We also qualitatively compared the frequency of mismatched individuals between brood (307 eggs and 213 larvae) produced by queens carrying at least a P haplotype and adults (workers, gynes, queens, and males) collected from polygyne colonies, We further investigated whether some types of mismatches are more common than others (P_2_-9a vs X-9r), and whether workers show higher mismatch frequencies than the reproductive caste.

Finally, we tested whether mismatch frequencies differ before and after mating in queens and males to indirectly assess whether observed genotype frequencies are likely to result from extrinsic selection (e.g., mating failure or unsuccessful colony founding, in which case we would expect a difference between gynes and mature queens or males and sperm) or from intrinsic genetic incompatibilities between haplotypes (in which case we would expect no difference in genotype frequency based on age).

We define *matched* individuals as those carrying the expected haplotype combination on each chromosome (P_2_-9r and X-9a, where X denotes any non-P_2_ haplotype on chromosome 3) (diploids: P_2_P_2_-9r9r, XP_2_-9a9r, XX-9a9a; haploids: P_2_-9r, X-9a). *Partially mismatched* individuals carry one matched (P_2_–9r or X–9a) and one mismatched (P_2_–9a or X–9r) haplotype combination and can only occur in diploid individuals. Within this category, we further distinguish individuals according to the matched haplotype combination they retain. We refer to individuals with one P_2_– 9r pair as *partial mismatches with a P*_*2*_*–9r pair* (P_2_P_2_–9a9r and XP_2_–9r9r) and those with one X– 9a pair as *partial mismatches with an X–9a pair* (XP_2_–9a9a and XX–9a9r). *Mismatched* individuals carry mismatch pairs on both chromosomes (diploids: P_2_P_2_-9a9a, XX-9r9r; haploids: X-9r, P_2_-9a) (see Table2).

**Table 2.** Definition of matched, partially mismatched, and mismatched genotypes. Genotypes are categorized according to the number of expected chromosome 3–chromosome 9 haplotype combinations (P_2_–9r and X–9a) they contain. Matched genotypes carry two expected pairs, partially mismatched genotypes carry one expected pair and one mismatched pair, and mismatched genotypes carry only mismatched pairs.

| **Genotype** | **P2-9r pairs** | **X-9a pairs** | **combination type** |
| --- | --- | --- | --- |
| P<sub>2</sub>P<sub>2</sub>-9r9r | 2 | 0 | matched |
| P<sub>2</sub>X-9a9r | 1 | 1 | matched |
| XX-9a9a | 0 | 2 | matched |
| P<sub>2</sub>P<sub>2</sub>-9a9r | 1 | 0 | Partial mismatched with a P<sub>2</sub>-9r pair |
| XP<sub>2</sub>-9r9r | 1 | 0 | Partial mismatched with a P<sub>2</sub>-9r pair |
| XP<sub>2</sub>-9a9a | 0 | 1 | Partial mismatched with a X-9a pair |
| XX-9a9r | 0 | 1 | Partial mismatched with a X-9a pair |
| P<sub>2</sub>P<sub>2</sub>-9a9a | 0 | 0 | mismatched |
| XX-9r9r | 0 | 0 | mismatched |

### Statistical analyses

To test whether haplotype transmission is biased by meiotic drive, we first analyzed eggs laid by heterozygous XP_2_ queens (chromosome 3) and 9a9r queens (chromosome 9). For each queen, we calculated the proportion of eggs inheriting the maternal P_2_ or 9r haplotype and fitted two generalized linear models (GLMs) with a binomial error distribution [using the glm function in R, lme4 package (Bates et al. 2015)], one for chromosome 3 and one for chromosome 9. In each model, the response variable was the proportion of eggs inheriting the maternal P_2_ (chromosome 3) or 9r (chromosome 9) haplotype, with male genotype (the queen’s mate) included as a fixed factor. Models were weighted by the total number of eggs analyzed per queen.

To test whether P_2_ and 9r haplotypes are transmitted independently and at equal frequencies consistent with Mendelian expectations (25% each), we modeled the maternal haplotype combination in eggs (categories P_2_-9a, P_2_-9r, P_1_-9a, P_2_-9r) as a multinomial response. The model included only an intercept as fixed effect and queen identity as a random intercept to account for repeated measures of eggs from the same queen. Deviation from the Mendelian expectation was assessed by inspecting whether the posterior predicted probabilities for each category overlapped 0.25. We excluded from this analysis double-mated queens. We used a Bayesian multinomial regression [brms, family = categorical, N = 79 eggs, 10 queens, 8000 post-warmup draws (Bürkner 2017)].

To test whether P_2_ and/or 9r act as maternal-effect killers to increase their representation in the next generation, we compared genotype frequencies in eggs and larvae produced by heterozygous P_1_P_2_ queens (chromosome 3) mated with P_1_ and M_A_ males and 9a9r queens (chromosome 9) mated with 9a males. For each queen, we calculated the proportion of P_2_ heterozygous and 9a9r offspring in eggs and larvae and fitted two generalized linear mixed models (GLMMs) with binomial error distributions (using the glm function in R), one for chromosome 3 and one for chromosome 9. In each model, the response variable was the proportion of offspring inheriting the maternal P_2_ (chromosome 3) or 9r (chromosome 9) haplotype, with developmental stage (eggs vs. larvae) included as a fixed factor and queen ID as a random effect. For chromosome 3, we additionally included mate genotype (P_1_ and M_A_) as a fixed factor and tested for its interaction with caste. Models were weighted by the total number of eggs and larvae analyzed for each proportion.

Deviation of maternal haplotype combinations (P_2_-9a, P_2_-9r, P_1_-9a, P_1_-9r) from the expected 25% in larvae produced by double heterozygous P_1_P_2_-9a9r mothers was analyzed with a Bayesian multinomial regression (brms, family = categorical, N = 139 (eggs= 79; larvae= 60), 12 queens, 8000 post-warmup draws) using the P_2_-9a haplotype combination as the reference category. Predictors included developmental stages (eggs and larvae) as a fixed effect and queen identity as a random intercept. We then refit the model with identical settings but using P_2_-9r as the reference category to facilitate interpretation of contrasts. We used the hypothesis function to test whether the transmission probability of each haplotype combination differed between developmental stages.

We next tested whether certain genotype combinations across chromosomes were overrepresented among workers, gynes, mated queens, their spermathecal content and males using Fisher’s exact test with Monte Carlo simulation (10,000 replicates) for each group (levels: 9a9a, 9a9r, 9r9r; P_2_P_2_, P_2_ heterozygous, non-P_2_ genotypes). For each group, we constructed contingency tables of genotype combinations and compared observed to expected counts under the assumption of independence between loci (i.e., no epistasis between the supergene on chromosome 3 and chromosome 9) using chi-square tests. Standardized residuals were calculated for each cell. For males, we only assessed haploid individuals, excluding two diploids. We tested for differences in mismatch frequencies between workers, gynes and mated queens using a Fisher’s exact test with Monte Carlo simulation (10,000 replicates; levels: matched, partially mismatched with a P_2_-9r pair, partially mismatched with a X-9a pair; we excluded the mismatched category because no individuals carried mismatches at both chromosome combinations).

Finally, to test whether mismatch frequencies in the adult population are driven by the failure of mismatched reproductive individuals to mate and reproduce, we compared match category distributions between mated (mature queens) and unmated (gynes) females using a Fisher’s exact test with Monte Carlo simulation (10,000 replicates; levels: levels: matched, partially mismatched with a P_2_-9r pair, partially mismatched with a X-9a pair; we excluded the mismatched category because no individuals carried mismatches at both chromosome combinations). Similarly, we compared the frequencies of matched and mismatched haplotypes between virgin males (before mating) and sperm recovered from spermathecae (after mating). However, because no mismatched haploid individuals were observed in either group, the contingency table contained no variation (only matched individuals), precluding a meaningful statistical comparison.

## Results

### No evidence for meiotic drive for either of the two supergenes

We fit two binomial GLMs to test whether maternal haplotype transmission on chromosomes 3 and 9 deviated from Mendelian expectations (50%). In P_1_P_2_ queens, transmission of P_2_ did not differ from 50% either when mated with M_A_ males (51.5% P_2_; Wald test: z = 0.15, p = 0.88), with P_1_ males (50.7% P_2_; Wald test: z = 0.08, p = 0.94), or with P_2_ males (59.5% P_2_; z = 1.39, p = 0.16; Fig. 1A). Similarly, we found no effect of mate genotype on transmission of 9r: 9a9r queens mated with 9a males (51.7% 9r; z = 0.36, p = 0.71) or 9r males (48.2% 9r; z = –0.26, p = 0.79) did not deviate from equal transmission (Fig. 1B).

**Figure 1.**
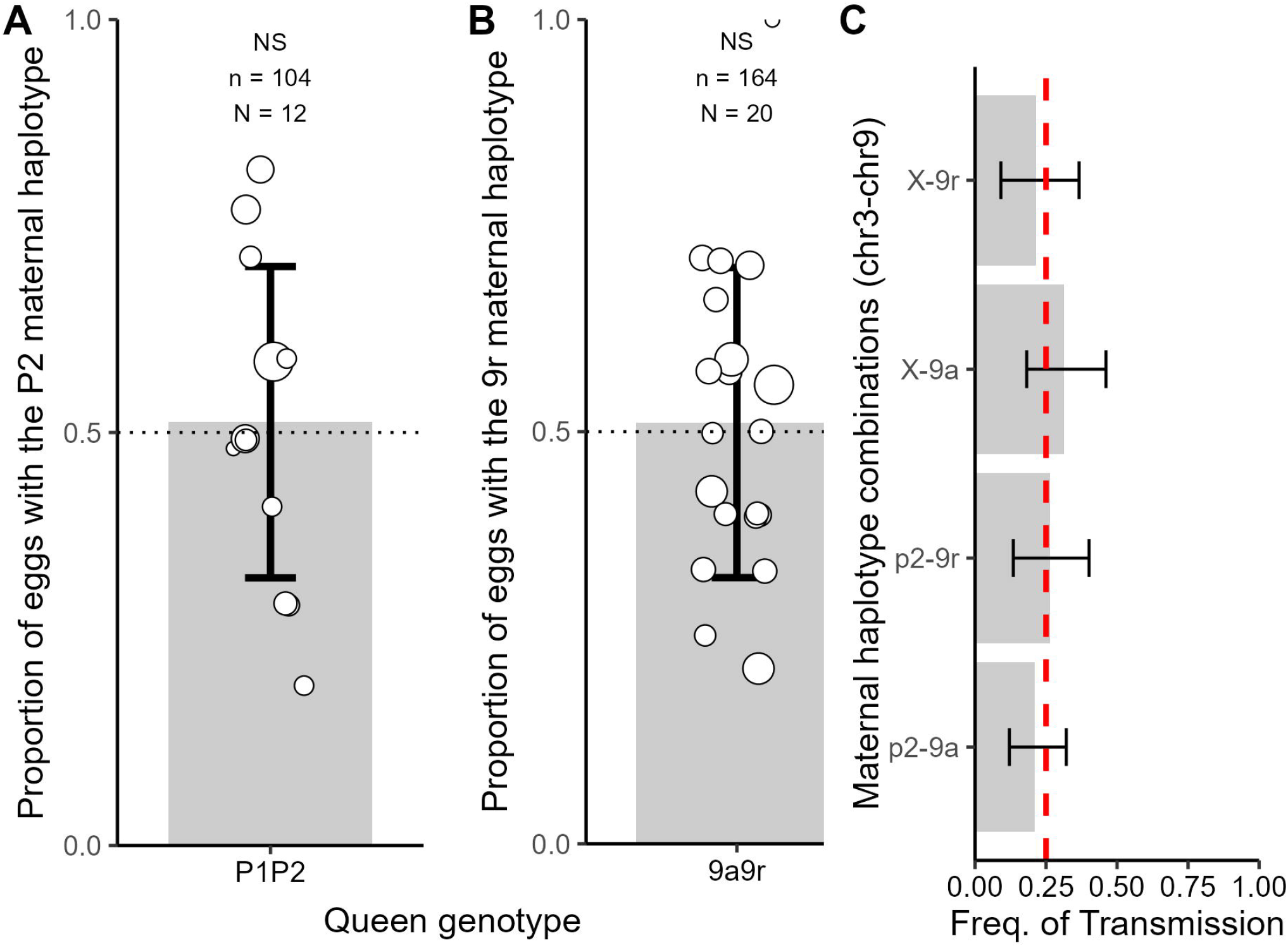
(A-B) Absence of meiotic drive: Maternal P_2_ and 9r haplotypes are transmitted to eggs in the expected 1:1 Mendelian ratio, indicating no evidence of meiotic drive. Error bars represent standard deviation, and dot size is proportional to the number of eggs analyzed per queen. (C) Independent assortment: Supergene haplotypes on chromosome 3 assort independently from those on chromosome 9, demonstrating that the elevated linkage disequilibrium (LD) between P_2_ and 9r is not caused by physical linkage. The barplot shows the multinomial regression model’s predicted mean frequency for each haplotype combination (averaged across all queens and mates), with error bars showing the 95% Bayesian credible intervals (CrI).

### Evidence of independent assortment between the supergenes

Across the 16 P_1_P_2_–9a9r queens, the four maternal haplotype combinations occurred at similar frequencies in eggs, averaging across all queens (P_1_–9r: 21.3%, 95% CrI = [9.3–35.5%]; P_1_– 9a:31.4%, 95% CrI = [18.7–46.2%]; P_2_–9r: 26.2%, 95% CrI = [13.9–39.9%]; P_2_–9a: 21.1%, 95% CrI = [12.4–39.9%]), with credible intervals overlapping the expected Mendelian frequency of 25% for all combinations. This reveals that haplotypes on chromosome 3 are transmitted independently from the haplotypes on chromosome 9 (Fig. 1C).

### No evidence that P_2_ or 9r are maternal-effect killers

To investigate whether P_2_ and/or 9r distort their transmission through maternal-effect killing, as discovered for the P haplotype in *F. selysi* (Avril et al. 2020), we fit generalized linear mixed models (GLMMs) to test for differences in the proportion of P_2_ heterozygous or 9a9r brood between the egg and larval stages. ANOVA tests revealed no significant difference in the proportion of heterozygous brood between eggs and larvae for either P_2_ (χ^2^ = 0.01, df = 1, p = 0.92, Fig. 2C) or 9r (χ^2^ = 0.05, df = 1, p = 0.82, Fig. 2B), suggesting that neither haplotype acts as a maternal killer with respect to P_1_ and 9a. For chromosome 3, we also included mate genotype (P_1_ and M_A_) as a fixed factor and tested its interaction with developmental stages; we found no significant effect of mate genotype on the proportion of P_2_ heterozygous brood (χ^2^ = 0.09, df = 1, p = 0.77), nor a significant interaction between developmental stages and mate genotype (χ^2^ = 0.01, df = 1, p = 0.90). To further test whether the transmission of the four haplotype combinations (P_2_-9a, P_2_-9r, P_1_-9a, P_1_-9r) differed from expected Mendelian ratios in larvae and whether their transmission differed compared to eggs, we fit a Bayesian multinomial regression model with queen as a random effect. Results for larvae mirrored those observed in eggs: posterior predictions showed similar transmission frequencies across all four haplotype combinations, averaging across all queens ( P_1_-9r: 22.2%, 95% CrI = [11.1%, 35.3%]; P_1_-9a: 29.6%, 95% CrI = [17.7%, 43%]; P_2_-9r: 31.3%, 95% CrI = [19%, 46.4%]; P_2_-9a: 16.9%, 95% CrI = [8.6%, 27.6%]), with all credible intervals overlapping the expected Mendelian frequency of 25%, indicating no evidence for transmission distortion (Fig. 2C). We did not detect any difference in the proportion of haplotype combinations between eggs and larvae (all caste effect credible intervals included zero: P_2_-9r caste estimate −0.41 [−1.52, 0.68]; P_2_-9a caste estimate= 0.31 [−0.72, 1.34]; P_1_-9a caste estimate= −0.18 [−1.23, 0.86]; P_1_-9r caste estimate= −0.23 [−1.38, 0.91]).

**Figure 2.**
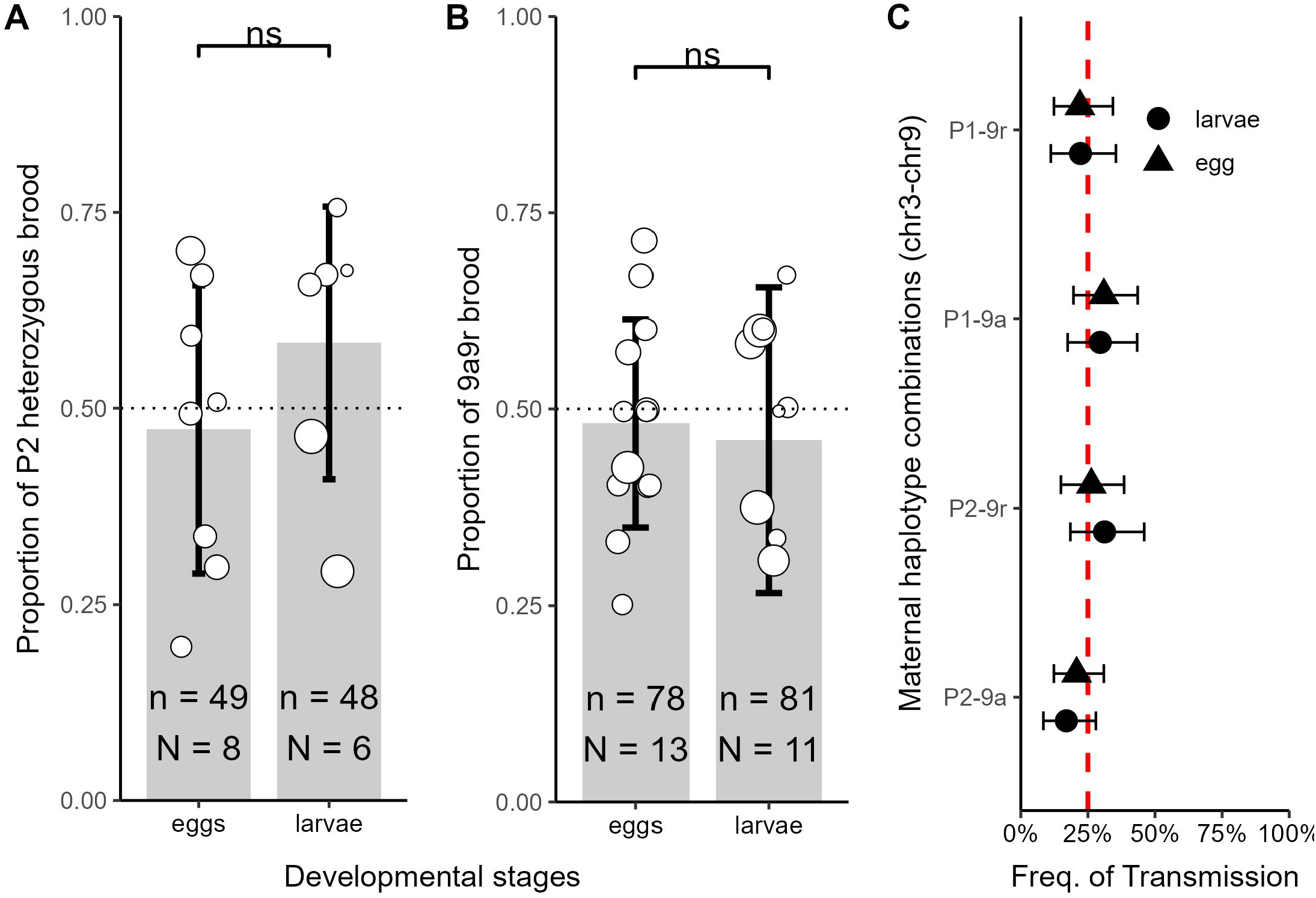
(A-B) Absence of maternal-effect killing: P_2_ and 9r do not act as maternal-effect killers, as the proportion of heterozygous brood does not differ between eggs and larvae. Error bars represent standard deviation, and dot size is proportional to the number of eggs analyzed per queen. (C) In larvae, the proportions of the four maternal haplotype combinations do not deviate from the expected 1:1:1:1 ratio, and do not differ from transmission observed in eggs, indicating that none of these haplotype combinations cause early mortality. The plot shows the multinomial regression model’s predicted mean frequency for each haplotype combination (averaged across all queens and mates), with error bars showing the 95% Bayesian credible intervals (CrI).

### Specific mismatched genotypes are underrepresented in adults

We next tested whether genotypes on chromosomes 3 and 9 were associated in workers, gynes, mated queens, spermathecae and males. Fisher’s exact tests revealed a highly significant association between the two supergenes in all three castes (workers: p < 0.0001; gynes: p < 0.0001; mated queens: p < 0.0001; spermathecae: p< 0.0001; males: p < 0.0001). Matched genotype combinations (with both chromosomes carrying matched haplotype pairs) were consistently overrepresented relative to expectations, partial mismatches (observable only in diploids individuals) with a P_2_-9r pair were found at the expected frequencies or overrepresented, whereas all partial mismatches with a X-9a pair were strongly underrepresented (Fig. 3A,B,C). Mismatched genotypes, where both chromosome pairs carried mismatched haplotype combinations (P_2_P_2_–9a9a and XX–9r9r) were completely absent (Fig. 3A). Likewise, none of the 39 spermathecal content and the 128 haploid males analyzed carried a mismatched genotype, except for a single diploid male with a partial mismatch (P_2_P_2_–9a9r) (Fig. 3B, Table S4). In broods (eggs and larvae) produced by queens carrying at least one P haplotype, mismatched individuals accounted for 3.7% of the offspring (n = 19), whereas partially mismatched individuals carrying an X–9a pair and those carrying a P_2_–9r pair accounted for 16.5% (n = 86) and 15.4% (n = 80), respectively (Tables S3-S4). Matched individuals represented 64.4% of the offspring (n = 335). In contrast, among adults from the polygyne population, we did not detect any mismatched individuals (Tables S3-S4). We identified 10 partially mismatched individuals carrying an X–9a pair (2.5%) and 29 partially mismatched individuals carrying a P_2_–9r pair (7.2%) among 404 diploid samples. Frequencies of matched and partially mismatched categories did not differ significantly among workers, gynes, and mated queens (Fisher’s exact test with Monte Carlo simulation, p = 0.1). We did not find a significant difference in the frequency of mismatches between unmated gynes and mated queens (p = 0.46). Finally, we did not find any mismatches in males or sperm.

**Figure 3.**
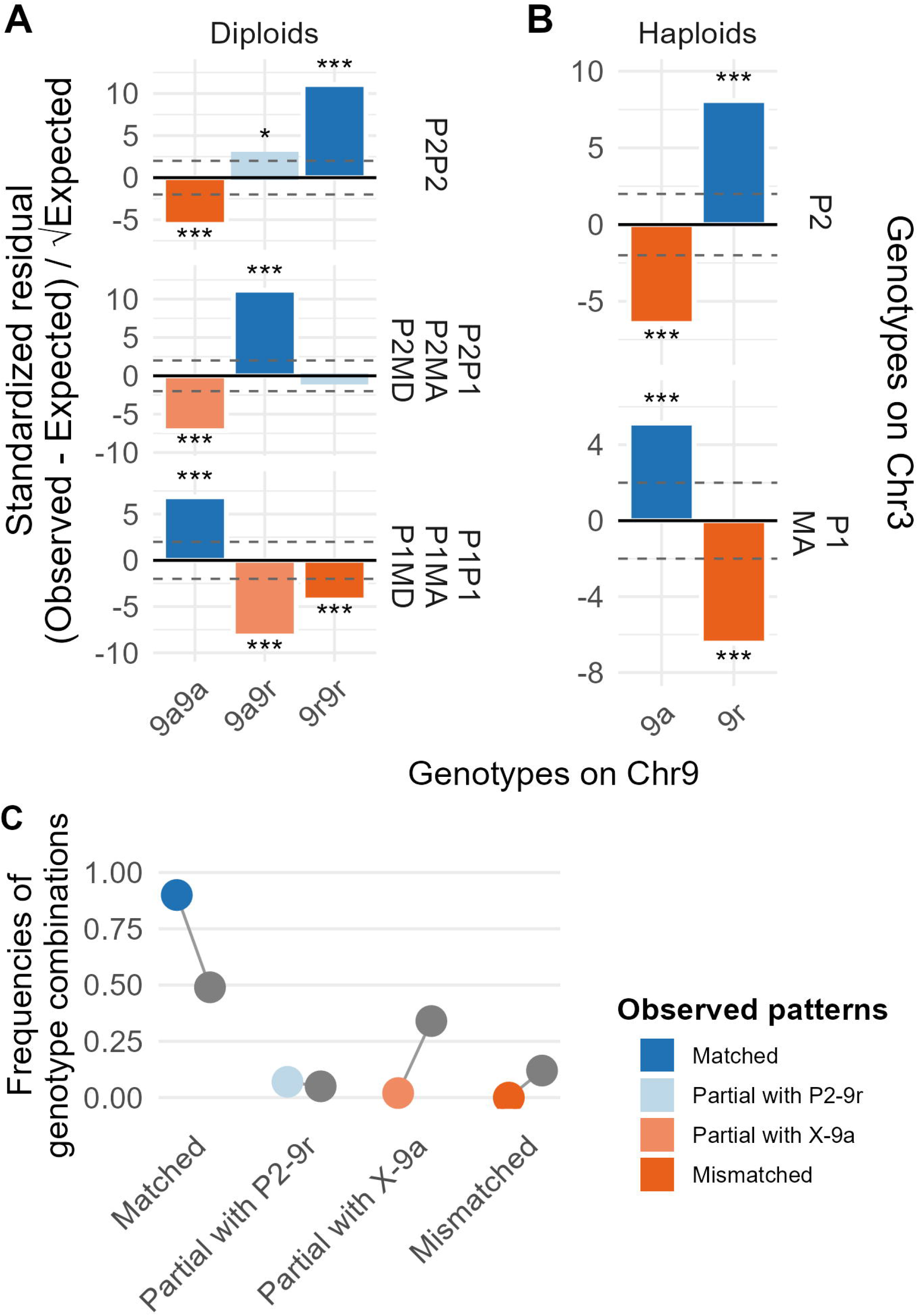
Comparison of expected and observed polygyne population-level genotype frequencies in diploid individuals analyzed with RAD-seq by Scarparo et al. (2023) and PCR-RFLP in this study(A): workers, gynes, and mated queens; and haploid individuals (B): spermathecal content and males. We only show the results for single-mated queens for the spermathecal content. (C) Expected (in gray) and observed mismatch frequencies among diploid polygyne individuals.

## Discussion

The formation of maladaptive allele combinations is often resolved through recombination suppression within supergenes (Thompson & Jiggins 2014). Here, we provide empirical evidence of strong epistasis between two supergenes in *F. cinerea* that control social organization (monogyne vs. polygyne) and queen and male size (macrogynes/macraners vs. microgynes/micraners). These supergenes exhibit strong linkage disequilibrium (LD), with the polygyne-associated P_2_ haplotype and the miniaturizing 9r haplotype co-occurring more frequently than expected (Scarparo et al. 2023). We demonstrate that, in this natural population, bidirectional epistatic interactions between two unlinked supergene regions impose severe viability costs on some mismatched genotype combinations, maintaining non-random haplotype associations across the genome without physical linkage and thereby coupling colony social form with queen body size.

### LD between supergenes is maintained by asymmetric selection against mismatched genotypes

Our study demonstrates that the P_2_ and 9r haplotypes assort independently during meiosis. Heterozygous P_1_P_2_–9a9r queens transmit all four possible haplotype combinations to eggs at the expected Mendelian ratio. This independent assortment rules out physical linkage as an explanation for the strong LD observed between supergenes. This finding is surprising because elevated LD typically results from reduced recombination between physically linked loci, which can occur due to close physical proximity on the same chromosomes or through chromosomal rearrangements, such as the inversions observed in many organisms [i.e. ruffs (Lamichhaney et al. 2016); fire ants (Yan et al. 2020); *F. selysi* (Brelsford et al. 2020); butterflies (Jay et al. 2021)], or through chromosomal fusion, as reported for mating-type loci in several fungi species (i.e. Duhamel et al. 2022; Lucotte et al. 2025).

Instead, in *F. cinerea*, the LD between supergenes appears to be maintained by epistatic interactions between haplotypes, as genotype frequencies in the adult population deviated significantly from expectations under independent assortment and from the observed genotype frequencies in eggs and larvae. The most striking evidence comes from haploid individuals: we found no males or sperm carrying a mismatched haplotype combination. Because males are haploid, they cannot compensate for a mismatch with a second matched copy, making this complete absence strong evidence that mismatched haplotype combinations are lethal in haploid individuals. Across diploids (workers, gynes, and queens), mismatches were also absent and partially mismatched combinations with a X-9a pair were severely underrepresented, consistent with complete or near-complete lethality. In contrast, partially mismatched genotypes with a P_2_-9r pair (P_2_P_2_-9a9r and XP_2_-9r9r) occurred at expected (or above expected) frequencies. Matched genotypes were also highly overrepresented. This pattern indicates that diploids carrying at least one P_2_ haplotype generally require at least one 9r haplotype to be viable. Together, these results are consistent with recessive epistatic lethality, where at least one matched P_2_-9r combination enables normal development.

Segregation distorters could account for deviations from expected genotype frequencies. We genotyped eggs and larvae to determine whether meiotic drive or a maternal-effect killer biased genotype ratios at early stages of development. Our data are not consistent with either segregation distortion mechanism. First, strong meiotic drive can be excluded: both P_2_ and 9r are transmitted to eggs at approximately 50% frequency by heterozygous queens, with no evidence that mate genotype influences transmission (consistent with normal Mendelian segregation). Second, a strong maternal-effect killer mechanism, analogous to the system in *F. selysi* (Avril et al. 2020), seems unlikely. Indeed, crosses between P_1_P_2_ queens and P_1_ or M_A_ males yield P_2_ heterozygous genotypes at the expected ~50% frequency in both eggs and larvae, and 9a9r queens mated to 9a males similarly produce expected genotype ratios in larvae. These results are not consistent with maternal-effect killers targeting specific genotype combinations, at least when considering transmission on P_2_ relative to P_1_ and 9r relative to 9a.

Assortative mating with regard to supergene haplotype has been reported in *F. selysi* (Avril et al. 2019) and for the Gp9 gene (later determined to be within the queen number supergene) in *S. invicta* (Ross & Keller 1995) and could theoretically produce some of the observed genotype frequency patterns in adults. However, a recent study on *F. cinerea* shows that while monogyne and polygyne macrogynes (without P_2_ and 9r) (see Table 1) mate assortatively with respect to their supergene genotype, heterozygous P_2_-9r microgynes mate randomly (Scarparo et al. 2026), ruling out assortative mating as an explanation for the observed genotype combination frequencies. Moreover, assortative mating alone is insufficient to explain several key observations, including the complete absence of mismatched males in our population samples, and the specific asymmetries in which mismatch types survive (see Fig. 3).

Taken together, our results point to fitness epistasis between the two supergenes as the primary mechanism shaping genotype frequencies in adults and elevated LD between chromosomes.

### Postzygotic incompatibilities couple social organization and body size

The chromosome 3 supergene controls social organization, with different haplotypes associated with polygyny (P_1_, P_2_) or monogyny (M_A_, M_D_). The chromosome 9 supergene controls the body size of queens and males, with tiny individuals having at least one copy of the 9r haplotype. In natural populations, these traits show strong phenotypic associations: the P_2_ and 9r haplotypes are found together far more frequently than expected by chance, with only polygyne colonies (and never monogyne colonies) containing tiny P_2_-9r reproductive individuals (Scarparo et al. 2023). This haplotype coupling could result from low fitness in adults carrying a mismatched haplotype combination, from workers discriminating against mismatched eggs and/or larvae during development, or intrinsic lethality in individuals with mismatched haplotype combinations. If extrinsic selection on adults drives the P_2_-9r association, mismatched males and gynes should develop normally and either fail to mate or be selected against post-mating (e.g., through unsuccessful colony foundation). In contrast, intrinsic lethality or worker-mediated selection would prevent mismatched individuals from completing development. We tested these alternatives by comparing mismatch frequencies across developmental stages (eggs, larvae, and adults) and between reproductives before and after mating (unmated gynes vs. mated queens and unmated males vs sperm in the female reproductive tract).

Across eggs and larvae (n = 520) produced by polygyne queens carrying at least one P haplotype (either P_1_ or P_2_), fully mismatched genotypes occurred at a frequency of 3.7%, while partially mismatched combinations carrying an X–9a pair occurred at a frequency of 16.5%. In contrast, among adults sampled from the entire polygyne population, fully mismatched genotypes were absent, and partially mismatched combinations carrying an X–9a pair were detected at only 2.5% frequency across queens, gynes, and workers. Similarly, of the 135 haploid males and 40 spermathecae we successfully dissected and genotyped, none carried mismatched genotype combinations (Fig. 3B, Table S4). All males were collected from colonies, some as pupae and some as newly eclosed adults, likely before mating (though we cannot exclude that some individuals mated with gynes within their natal colony). The complete absence of mismatched genotypes in haploids before (the males) and after (the sperm) mating strongly suggests that mismatched haplotype combinations are purged during development rather than causing reduced fecundity or mating success. Indeed, if mismatched males developed normally but failed to mate, we would expect to observe them among the unmated male samples. Consistent with this interpretation, we detected no significant differences in mismatch frequency between unmated gynes and mated queens, further supporting developmental viability selection over mating failure. Our dataset does not allow us to clearly distinguish between intrinsic lethality and worker-mediated selection, as all broods were reared in the presence of workers. Nevertheless, worker discrimination seems unlikely for several reasons. First, only certain mismatch classes are underrepresented, while others occur at expected frequencies (Figure 3). This pattern would require extremely fine-grained discrimination given the genotypic complexity of this two-supergene system. If workers were responsible, CHC-based recognition would be the most plausible mechanism, as in *Solenopsis invicta*, where polygyne workers selectively kill queens lacking the Sb haplotype (Keller & Ross 1998; Trible & Ross 2016; Zeng et al. 2022); yet even there, discrimination appears limited to queens. Consistent with this, MM and MP workers of *Formica selysi* show similar CHC profiles (Choppin et al. 2026), arguing against supergene-mediated worker-worker discrimination. By analogy, if *F. cinerea* workers could recognize haplotype combinations, we would predict higher mismatch frequencies among workers than among gynes and mated queens, a pattern we do not observe. Second, while ants readily discriminate non-nestmate adults, *Formica* as a genus shows weak or no discrimination against immature stages (Schultner & Pulliainen 2020), further arguing against worker-mediated removal of mismatched brood. Third, the complete absence of mismatched haploid individuals is difficult to link with worker-mediated selection: workers would have to be imperfect selectors, as some diploid mismatches survive, making it implausible that they would eliminate every single haploid mismatch. This asymmetry is more consistent with intrinsic lethality, where the absence of a second matched copy drives haploid mismatches to lethality. We therefore propose that intrinsic lethality during development is the most parsimonious explanation for the paucity of mismatched adults and for the maintenance of both the genetic (P_2_–9r) and phenotypic (polygyne–small body size) associations.

We acknowledge that our approach, based on eliminating alternative mechanisms rather than directly observing the cause of mortality, does not allow us to identify the precise developmental mechanism responsible for removing mismatched genotypes. However, we rule out physical linkage, meiotic drive, maternal-effect killing, and assortative mating as explanations for the observed linkage disequilibrium. We also exclude the possibility that P_2_-9r coupling results from mating or post-mating extrinsic selection. Consequently, the strong depletion of partially mismatched combinations carrying an X–9a pair, together with the complete absence of fully mismatched individuals among adults, most likely reflects genotype-dependent postzygotic viability selection. Whether this selection reflects intrinsic genetic incompatibilities between the two supergenes or is mediated by worker selection does not change the evolutionary inference: survival depends on the interaction between genotypes at the two supergene regions, which by definition constitutes fitness epistasis.

### Epistatic incompatibility between supergenes mirrors a synthetic gene drive

The severity of epistatic interactions depends critically on mating combinations. Queens carrying P_2_–9r haplotypes and heterozygous at one or both supergene regions experience dramatically different offspring viability depending on the mate genotype (Table 3). When mated with P_1_–9a or M_A_-9a males, these queens are expected to produce many lethal genotype combinations, resulting in up to 75% offspring mortality. In contrast, mating with P_2_–9r males can rescue offspring that received a mismatched maternal haplotype combination, yielding predominantly viable offspring. Because mismatched males never survive to reproduce, mismatched haplotype combinations can only enter the next generation through the queens. This system strikingly mirrors the synthetic threshold-dependent maternal-effect lethal underdominance (UD^MEL^) engineered in *Drosophila* by Akbari et al. (2013), in which two maternally expressed toxins on separate chromosomes are each linked to a zygotic antidote rescuing the lethality induced by the other toxin. Individuals that fail to inherit at least one matched toxin–antidote pair do not survive. Analogously, heterozygous females carrying one UD^MEL^ chromosome mated to wildtype males suffer severe fitness costs, with offspring dying as embryos or first-instar larvae, closely paralleling the inferred fitness cost of heterozygous P_2_-9r queens mated with P_1_-9a or M_A_-9a males.

**Table 3.** Offspring viability depends on queen-male genotype combinations. Queens carrying P_*2*_-9r haplotypes produce predominantly viable diploid offspring when mated with P_*2*_-9r males, but largely unviable offspring when mated with X-9a males.

| Parental genotypes | P<sub>2</sub>-9r |  | X-9a |  |
| --- | --- | --- | --- | --- |
| P<sub>2</sub>P<sub>2</sub>-9r9r | P<sub>2</sub>P<sub>2</sub>-9r9r |  | P<sub>2</sub>X-9a9r |  |
| P<sub>2</sub>P<sub>2</sub>-9a9r | P<sub>2</sub>P<sub>2</sub>-9r9r | P<sub>2</sub>P<sub>2</sub>-9a9r | P<sub>2</sub>X-9a9r | P<sub>2</sub>X-9a9a |
| P<sub>2</sub>X-9a9r | P<sub>2</sub>P<sub>2</sub>-9r9r | P<sub>2</sub>P<sub>2</sub>-9a9r | P<sub>2</sub>X-9a9r | P<sub>2</sub>X-9a9a |
|  | P<sub>2</sub>X-9a9r | P<sub>2</sub>X-9r9r | XX-9a9a | XX-9a9r |
| P<sub>2</sub>X-9r9r | P<sub>2</sub>P<sub>2</sub>-9r9r | P<sub>2</sub>X-9r9r | P<sub>2</sub>X-9a9r | XX-9a9r |
| XX-9a9a | P<sub>2</sub>X-9a9r |  | XX-9a9a |  |

However, key differences with the UD^MEL^ system exist. While *Drosophila* lethality in the UD^MEL^ system arises early in development, as also shown for the maternal-effect killer in *F. selysi* (Avril et al. 2020), our data suggest that lethality in *F. cinerea* occurs during late larval or pupal stages. The supposedly late mortality makes it unlikely that lethality of mismatched individuals in *F. cinerea* is driven by maternally (and paternally) inherited toxin–antidote interactions. Notably, P_1_ is also a polygyne haplotype on chromosome 3 yet does not show tight LD with 9r, indicating that whatever drove the co-evolution of P_2_ and 9r is specific to P_2_ and not a general property of polygyne haplotypes. However, what molecular mechanism underlies this naturally evolved incompatibility system remains an open and intriguing question.

## Conclusion

The *F. cinerea* system reveals an unusual mechanism by which linkage disequilibrium between unlinked genomic regions can be generated and maintained without requiring physical linkage. Instead, LD is maintained by epistatic selection against mismatched haplotype combinations, with mismatched individuals failing to survive until adulthood. This results in functional coupling of social organization and body size. Strikingly, the population-level dynamics of this naturally evolved system mirror the two-locus synthetic gene drive deliberately engineered in the laboratory in *Drosophila* (Akbari et al. 2013). The mechanistic basis of these epistatic interactions remains an important avenue for future research. Understanding why specific mismatched combinations show complete lethality (P_2_-9a and X-9r in haploid males) while others show reduced viability could reveal the functional relationships between these supergene regions. More broadly, this system highlights that complex adaptive phenotypes may not always require physical linkage to be coupled. Interactions among supergenes may represent a previously unexplored route by which co-adapted trait combinations can evolve and persist. As supergenes are increasingly recognized across diverse taxa, including emerging examples involving multiple supergene regions, an important future challenge will be to determine how frequently complex phenotypes are shaped and maintained across generations by epistatic interactions within and between supergenes.

## Acknowledgments

We thank Marie Palanchon and Marco Molfini for their assistance in collecting *F. cinerea* queens, and Zul Alam, Tom Nonacs, and Samantha Sudoko for feedback on the manuscript. This work was supported by the US National Science Foundation Division of Environmental Biology grant #1754834 to AB and JP and grant #1942252 to JP.

## Data, Materials, and Software Availability

All data generated in this study are available in the Supplementary Materials. Primer sequences are provided in Table S1. Complete genotyping data for chromosomes 3 and 9 are provided in Table S2, including data from individuals not included in the present study because they were produced by queens with non-target genotypes (e.g., MM or 9a9a). Genotyping data for the individuals included in the analyses presented in this study are provided in Table S3.

