## Supplementary material for "Lethal epistasis maintains strong linkage disequilibrium between unlinked supergenes": Table S1, Table S4, Figure S1, Figure S2

### Supplementary files

#### PCR-RFLP

##### *DNA Extraction*

We extracted DNA from eggs, larvae, queens, and the queens' spermathecae. For eggs and larvae, individual samples were placed into wells of a 96-well plate and digested overnight at 57 °C in a mixture of Buffer ATL and Proteinase K. The following day, the entire supernatant was transferred to a deep-well plate for automated extraction using the QIAcube HT/QIAxtractor system, using the QiaAmp 96 protocol for tissue. For queen spermathecae, we used the same protocol, but queens preserved in ethanol were first dissected under a Leica S8APO microscope. Intact spermathecae were then transferred into wells of a 96-well plate and digested overnight in Buffer ATL and Proteinase K. For queens, the protocol was the same, except that tissues were first ground with pestles following immersion in liquid nitrogen before overnight digestion in Buffer ATL and Proteinase K. DNA from all samples was eluted in 100 µL of EB Buffer.

##### *DNA amplification and enzymatic digestion*

We designed seven PCR-restriction fragment length polymorphism (PCR-RFLP) assays for haplotype identification in *F. cinerea*: five assays to genotype the four supergene haplotypes on chromosome 3 and two assays to genotype the two haplotypes on chromosome 9. Given the low DNA quantities typically obtained from single eggs or spermathecae, we implemented a nested PCR strategy using two sequential amplification steps with external and internal primer pairs (Table S1). Haplotype identification was achieved by pairing each internal primer set with a restriction enzyme that recognizes a diagnostic single-nucleotide polymorphism (SNP), producing distinct restriction fragment patterns for each haplotype.

**Table S1**. Overview of primers and associated restriction enzymes.

| # | External Primer Sequences | Product Size | Internal Primer Sequences | Product Size | Enzyme | Enzyme working Temperature | Size of digested Product | Targeted Haplotype | Chromosome | Comments |
|---|---|---|---|---|---|---|---|---|---|---|
| **1** | TTCTGCGCTATATGTTATGCCTAT<br>TGGGACGAAACGATTTTCA | 343 bp | TGTTcATAAGAAAATGATACCTGGA<br>GGCGGTGCCAATCAACTAT | 170 bp | BtscI | 50 °C | 104 bp<br>66 bp | P2 | 3 | BtscI leaves P2 intact; digested DNA segments can be MA, MD, P1 |
| **2** | AAACGATCCGTCCGTCTTC<br>ATAAAGTCCGCTCGCTGAGA | 240 bp | CGAGATCTGCAGGGCTCTACT<br>AGAGCCGCGACCAAGTTATT | 156 bp | RsaI | 37 °C | 98 bp<br>58 bp | P1 | 3 | RsaI leaves P1 intact; digested DNA segments can be MA, MD, P1 |
| **3** | AGGGGGAAGGGAAAGAGAAT<br>GAATGCGAATACGGAGCAGT | 245 bp | GACAGAGTGCAGCAATTTCG<br>GTCGCGAAAGGAGAAGACTG | 163 bp | Sau96I | 37 °C | 90 bp<br>73 bp | P1 | 3 | Sau96I cuts P1 and leave MA, MD, P2 undigested |
| **4** | CGTTTCACGCGAATTTGTC<br>TCCCGGCTGTCTCGTTAAT | 161 bp | AATTTGTCGCGCACATCG<br>GGCTGCAGGGCTTGTTAGT | 150 bp | Hpy166II | 37 °C | 87 bp<br>63 bp | MD | 3 | Hpy166II leaves MD intact; digested DNA segments can be MA, P1,P2 |
| **5** | CGCTTTCCTCGTGCTAAAGT<br>TGGAACTCATCCCCTGCTAC | 247 bp | GTGCTAAAGTCCAGCACAGC<br>CCTGCTACCACCCCATTCT | 225 bp | HpaII | 37 °C | 147 bp<br>47 bp | MP | 3 | HpaII cuts the M haplptypes (MA and MD) and leaves intact the P hapltypes (P1 and P2) |
| **6** | GCGACTCGTGCATGATCTG<br>CAATACGCCGAACCTGCTC | 295 bp | CCCATACTCGCTTGCATGC<br>CTCGCTTTGCCTCCCTTTG | 197 bp | TaqI-v2 | 65 °C | 101 bp<br>96 bp | 9a | 9 | TaqI-v2 cuts 9a and leaves 9r intact |
| **7** | GGCAAAAGCGTACTACATCG<br>TCGTCGCCCATATTCTTCGA | 319 bp | TGCACTTTACGACACTGGC<br>AGATTCGGCATTGGGTCTCT | 186 bp | Hpy166II | 37 °C | 117 bp<br>69 bp | 9r | 9 | Hpy166II cuts 9r and laeves 9a intact |

Each 10 µl PCR reaction included 6.8 µl $ddH_2O$, 1 µl of 10 µM pre-mixed forward and reverse primers, 1 µl ThermoPol buffer, 0.08 µl Taq polymerase, 0.08 µl of 25 mM dNTPs, and 1 µl DNA template. The PCR program consisted of an initial denaturation at 94 °C for 3 min, followed by 35 cycles of 94 °C for 30 s, 58 °C for 30 s, and 72 °C for 30 s; and a final elongation at 72 °C for 3 min. After the first round of amplification with the external primers, PCR products were diluted 1:10 in $ddH_2O$ and used as templates for a second PCR with the internal primers, using the same reaction mix and cycling conditions.

Each final PCR product was digested for 1 hour at the enzyme's recommended temperature using the following 9 µl reaction: 5.3 µl $ddH_2O$, 0.6 µl CutSmart buffer, 0.1 µl restriction enzyme, and 3 µl PCR product. The entire digest was then loaded on a 2% agarose gel and run for 55 minutes alongside a 100 bp DNA ladder.
Primers were designed using Primer3, and enzymes selected using NEBcutter v3.0.

*Chromosome 3 genotyping and dataset comparison*

To establish the most reliable threshold for genotype calling while minimizing sample exclusion, we generated two datasets with different stringency criteria. In the strict dataset, genotypes were called only when all assays produced concordant results; individuals with at least one discordant or failed assay (returning "NA") were classified as unknown ("unkn"). In the loose dataset, the "unkn" classification was assigned only when at least two assays were discordant, or when one was discordant and another failed. For both datasets, samples were classified as "NA" when at least two assays failed to yield readable results.
Of 1,662 individuals genotyped (see Table S2), the strict dataset successfully genotyped 1,397 individuals (158 unkn, 118 NA), while the loose dataset successfully genotyped 1,494 individuals (61 unkn, 107 NA).
To determine which dataset to use for downstream analyses, we tested whether they yielded consistent results for meiotic drive and maternal effect killing (see Meiotic Drive and Maternal Effect Killing sections in the main text). For meiotic drive, we fit a binomial GLMM with the proportion of maternally transmitted $P_2$ haplotype, when queens always carry the $P_1P_2$ genotype, as the response variable, dataset type as a fixed effect, queen ID as a random effect, and weighted by the total number of eggs per queen. The two datasets did not differ significantly ($z = 0.016$, $p = 0.98$; Fig. S1A). For maternal effect killing, we fit a similar model including caste as an additional fixed effect with its interaction with dataset type. Pairwise comparisons using emmeans revealed no significant differences between datasets or castes (all $p > 0.05$; Figure S1B). Given this concordance, we used the loose dataset for all subsequent analyses to maximize sample retention while maintaining reliability.

*Chromosome 9 genotyping*

For chromosome 9, we developed two assays (Table S1) and genotyped 1,253 individuals (see Table S2): 1,113 with concordant results from both assays, and 407 using a single assay (genotypes called only when results were unambiguous, e.g., bright bands). Forty-two samples were classified as "unkn" due to assay failure or discordant results.
For both chromosomes, the majority of unknown genotypes originated from eggs, larvae, and spermathecae (94.9% for chromosome 3; 85.7% for chromosome 9).

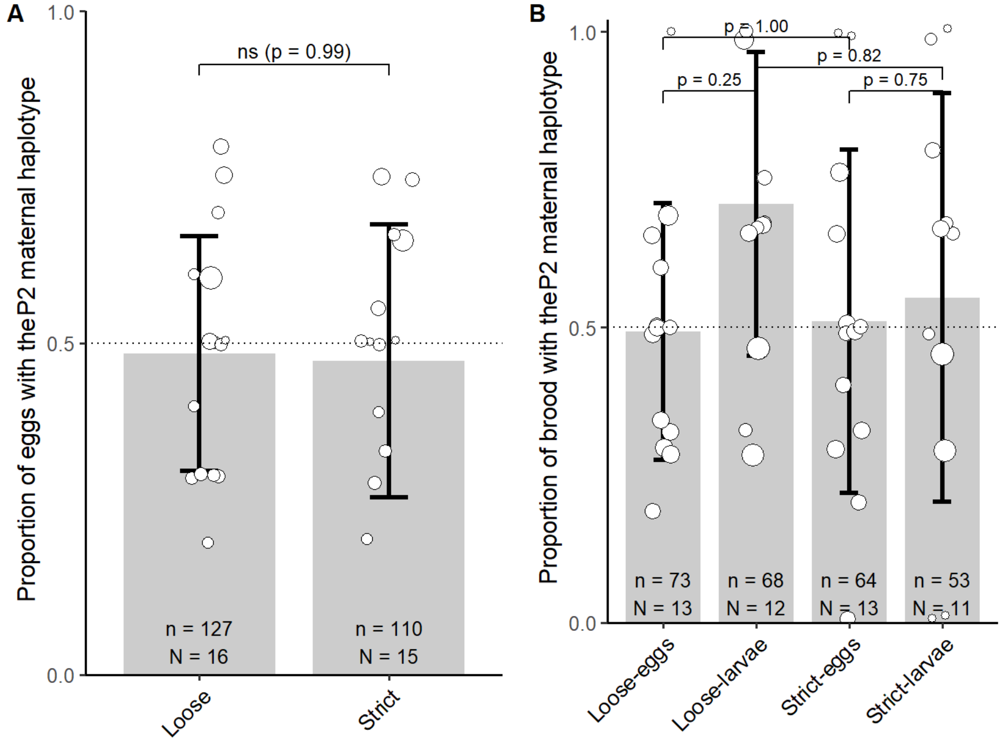


**Figure S1**. Comparison of strict and loose datasets for chromosome 3 analyses. (A) Meiotic drive: proportion of maternally transmitted P2 haplotype did not differ between datasets ($z = 0.017$, $p = 0.99$). (B) Maternal effect killing: proportion of maternally transmitted $P_2$ haplotype across castes showed no significant differences between datasets (all $p > 0.05$). Error bars represent ± SD.

**Weak or absent maternal-effect killing**

Our data show that $P_2$ does not act as a maternal-effect killer as demonstrated in Formica selysi, where MM eggs laid by MP queens mated to M males fail to hatch (Avril et al. 2020). However, in *F. selysi*, enhanced transmission of the maternally transmitted P haplotype was observed compared to the M haplotype, whereas in this study the $P_2$ haplotype is compared to $M_A$, $M_D$, and $P_1$ haplotypes combined (collectively termed X) due to insufficient replicates. In our dataset, we only have brood data for 4 $MP_2$ queens; however, we still find MM larvae, despite an increased frequency of the $P_2$ haplotype from eggs to larvae ($z = -3.073$, $p = 0.01$). This suggests that if the $P_2$ haplotype enhances its transmission over M haplotypes through maternal effect killing, this drive is weaker than in *F. selysi*, where 100% of larvae carry the P haplotype.

In contrast, the $P_2$ haplotype compared to $P_1$ maintains its frequency at approximately 50% across both eggs and larvae.

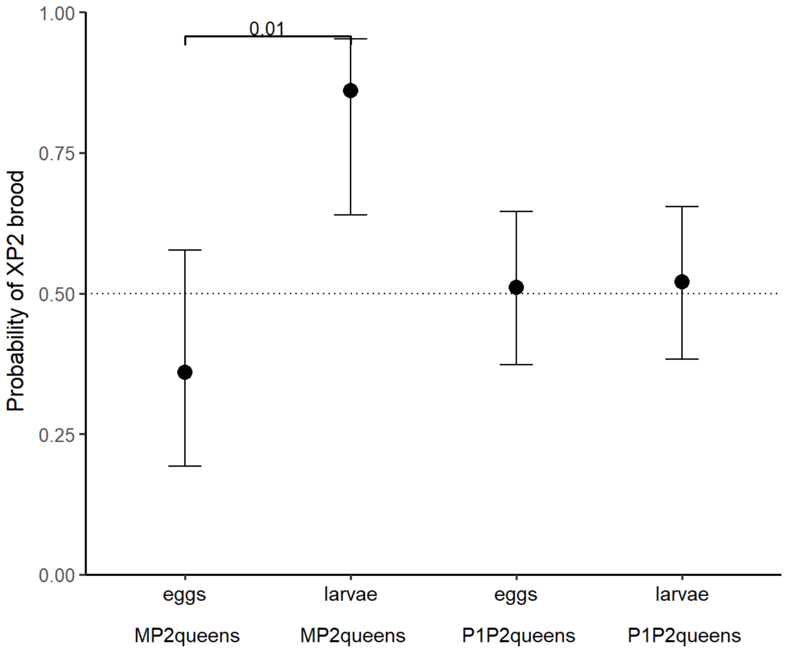


**Figure S2.** Details of maternal effect killing. The increase in the proportion of the $P_2$ haplotype was tested by fitting a GLMM with the proportion of the maternally transmitted $P_2$ haplotype as the response variable, caste and queen genotype ($MP_2$ or $P_1P_2$), and their interaction as fixed effects, queen ID as a random effect, and weighted by the total number of brood per queen. The $P_2$ haplotype shows an increased frequency in larvae produced by $MP_2$ queens; however, the skew is not 100%, as we still find MM larvae, suggesting that the *F. cinerea* system differs from the *F. selysi* gene drive system. In $P_1P_2$ queens, the transmission of $P_2$ in eggs and larvae remains invariant at 50%. Error bars represent 95% confidence intervals.

**Table S2.** Complete dataset of all individuals collected and genotyped using PCR–RFLP. The table includes both individuals used in this study and individuals produced by queens carrying uninformative genotype combinations (e.g., MM-9a9a), which were not included in the analyses presented here.

**Table S3.** Dataset of all individuals used in this study. This dataset includes individuals genotyped using PCR–RFLP and individuals genotyped using RAD-seq by Scarparo et al. (2023).

**Table S4**. Overview of genotypes across castes and developmental stages of samples analyzed in this study and Scarparo et al. 2023. We found a few eggs and larvae carrying mismatches in both chromosome pairs. Based on the genotypes of other eggs and larvae produced by the same queens and the genotype of the mate, we inferred that these individuals are haploid. The table shows only individuals successfully genotyped on both supergenes.

| Genotypes | | This study | | | | Scarparo et al. 2023 | | | |
|---|---|---|---|---|---|---|---|---|---|
| **Chr3** | **Chr9** | **eggs** | **larvae** | **spermathecae** | **queens** | **workers** | **gynes** | **males** | **queens** |
| $M_AM_A$ | 9a9a | - | 1 | - | - | 2 | - | - | - |
| $M_AM_A$ | 9a9r | - | 2 | | - | - | - | - | - |
| $M_AM_D$ | 9a9a | 4 | 1 | - | - | 3 | - | - | - |
| $M_AM_D$ | 9a9r | 4 | 1 | - | - | - | - | - | - |
| $P_1M_A$ | 9a9a | 13 | 17 | - | - | 155 | 16 | - | 5 |
| $P_1M_A$ | 9a9r | 6 | 8 | - | - | 1 | 1 | | - |
| $P_1M_D$ | 9a9a | - | 3 | - | - | 25 | 6 | 1 | 1 |
| $P_1P_1$ | 9a9a | 37 | 24 | - | - | 18 | 21 | - | 7 |
| $P_1P_1$ | 9a9r | 3 | 5 | - | 1 | - | | - | 1 |
| $P_1P_1$ | 9r9r | 8 | - | - | - | - | - | - | - |
| $P_1P_2$ | 9a9a | 12 | 8 | - | - | 3 | - | - | - |
| $P_1P_2$ | 9a9r | 33 | 21 | - | 11 | 27 | 11 | - | 8 |
| $P_1P_2$ | 9r9r | 19 | 16 | - | 3 | 2 | | - | - |
| $P_2M_A$ | 9a9a | 13 | 24 | - | - | 4 | - | - | - |
| $P_2M_A$ | 9a9r | 33 | 30 | - | 5 | 10 | 3 | - | 1 |
| $P_2M_A$ | 9r9r | 5 | 3 | - | - | - | - | - | - |
| $P_2M_D$ | 9a9r | - | - | - | 3 | - | 9 | - | - |
| $P_2P_2$ | 9a9a | 5 | 6 | - | - | - | - | - | - |
| $P_2P_2$ | 9a9r | 25 | 12 | - | 6 | 11 | 7 | 1 | - |
| $P_2P_2$ | 9r9r | 87 | 31 | - | 10 | 6 | 7 | - | 1 |
| $M_A$ | 9a | - | - | 15 | - | - | - | 7 | - |
| $M_D$ | 9a | - | - | - | - | - | - | - | - |
| $P_1$ | 9a | - | - | 13 | - | - | - | 72 | - |
| $P_2$ | 9r | - | - | 12 | - | - | - | 56 | - |
